# Surface-aerosol stability and pathogenicity of diverse MERS-CoV strains from 2012 - 2018

**DOI:** 10.1101/2021.02.11.429193

**Authors:** Neeltje van Doremalen, Michael Letko, Robert J. Fischer, Trenton Bushmaker, Claude Kwe Yinda, Jonathan Schulz, Stephanie N. Seifert, Nam Joong Kim, Maged G Hemida, Ghazi Kayali, Wan Beom Park, Ranawaka APM Perera, Azaibi Tamin, Natalie J. Thornburg, Suxiang Tong, Krista Queen, Maria D. van Kerkhove, Young Ki Choi, Myoung-don Oh, Abdullah M. Assiri, Malik Peiris, Susan I. Gerber, Vincent J. Munster

**Affiliations:** Laboratory of Virology, Division of Intramural Research, National Institute of Allergy and Infectious Diseases, National Institutes of Health, Hamilton, MT, 59840, USA; Department of Internal Medicine, Seoul National University College of Medicine, Seoul, South Korea; Department of Microbiology, College of Veterinary Medicine, King Faisal University, Al-Hasa, Saudi Arabia; Department of Virology, Faculty of Veterinary Medicine, Kafrelsheikh University, Kafrelsheikh, Egypt; Department of Epidemiology, Human Genetics, and Environmental Sciences, University of Texas Health Sciences Center, Department of Epidemiology, Human Genetics, and Environmental Sciences, Houston, Texas; School of Public Health, University of Hong-Kong, Hong Kong SAR, China; Division of Viral Diseases, National Center for Immunization and Respiratory Diseases, Centers for Disease Control and Prevention, Atlanta, GA, USA; Department of Infectious Hazards Management, Health Emergencies Programme, World Health Organization, Geneva, Switzerland; College of Medicine and Medical Research Institute, Chungbuk National University, Cheongju City, Republic of Korea; Infection Prevention and Control, Assistant Deputy Minister, Preventive Health, Ministry of Health, Riyadh, Saudi Arabia; Paul G. Allen School of Global Animal Health, Washington State University, Pullman, WA, 99111, USA

## Abstract

Middle East Respiratory Syndrome coronavirus (MERS-CoV) is a coronavirus that infects both humans and dromedary camels and is responsible for an ongoing outbreak of severe respiratory illness in humans in the Middle East. While some mutations found in camel-derived MERS-CoV strains have been characterized, the majority of natural variation found across MERS-CoV isolates remains unstudied. Here we report on the environmental stability, replication kinetics and pathogenicity of several diverse isolates of MERS-CoV as well as SARS-CoV-2 to serve as a basis of comparison with other stability studies. While most of the MERS-CoV isolates exhibited similar stability and pathogenicity in our experiments, the camel derived isolate, C/KSA/13, exhibited reduced surface stability while another camel isolate, C/BF/15, had reduced pathogenicity in a small animal model. These results suggest that while betacoronaviruses may have similar environmental stability profiles, individual variation can influence this phenotype, underscoring the importance of continual, global viral surveillance.

## Introduction

Middle East respiratory syndrome coronavirus (MERS-CoV) was first discovered in 2012 and continues to cause outbreaks in the Middle East as a result of frequent spillover from dromedary camels to humans. MERS-CoV has a mortality rate of ∼35% and has spread to 27 countries (1, 2). While dromedary camels have been shown to be the animal reservoir, phylogenetic analysis has shown that humans are a dead-end host (3). Approximately 41% of MERS-CoV cases in the Kingdom of Saudi Arabia (KSA) are primary, resulting from direct camel-to-human transmission (4). To date, MERS-CoV has been detected in camels in Burkina Faso, Egypt, Ethiopia, Jordan, Kenya, Morocco, Nigeria, Saudi Arabia, Senegal, Sudan, Tunisia, and Uganda (5-13). While MERS-CoV has been isolated from camels in Africa, there are no reports of zoonotic transmission to humans, unlike what has been observed in the Middle East (9).

Human-to-human transmission of MERS-CoV has been reported but is inefficient and primarily occurs in hospital settings and within households (14). The exact route of transmission between humans is currently still unclear. It is possible that direct contact with infected individuals, as well as fomite and aerosol transmission all collectively contribute to viral transmission. Epidemiological studies have mapped indirect patient contact within hospitals, providing evidence for aerosol and hospital-worker mediated spread (15-18). The largest outbreak of MERS-CoV outside of the Middle East occurred in South Korea. A single traveler from the Middle East brought MERS-CoV to South Korea, resulting in 185 subsequent infections (19).

Coronaviruses are the largest non-segmented RNA-based viruses identified, with genome sizes averaging around 30kB. Even though coronavirus polymerase has a proofreading function, coronaviruses within the same species are polymorphic. A recent study found approximately 99% nucleotide similarity as well as small deletions in nonstructural proteins between various isolates collected in the Middle East and North Africa (20). While this variation may seem minimal, 1% is equivalent to 300 nucleotide changes in the 30kB genome. Indeed, many of these changes are nonsynonymous and are distributed throughout the viral genome. As previously shown, single amino acid changes in the viral genome can result in profoundly varying phenotypes in viral replication (21). Additionally, one study has aimed to functionally characterize some of these MERS-CoV strain differences, with a particular focus on ORF deletions, and found significant effects in the virus’ ability to antagonize host innate immune pathways, translating to viral attenuation in an animal model (20). Given that MERS-CoV continues to cause outbreaks and evolve, these findings underscore the importance of characterizing how MERS-CoV genetic variation alters viral replication, pathogenicity and stability.

Here we expand on previous work from others by testing a broad panel of viral isolates collected from both humans and camels, representing every major geographic region with MERS-CoV outbreaks and spanning from early to contemporary outbreaks. Because MERS-CoV spreads within households and hospitals, we characterized and compared viral phenotypes with immediate implications for public health. We focused on environmental stability both in aerosols as well as surface stability on common materials found in hospitals, replication kinetics in immortalized human cell lines and primary human airway epithelial cultures, as well as pathogenicity in a transgenic mouse model our lab previously developed to test vaccine efficacy (22). For the environmental stability studies, we included SARS-CoV-2 to allow better comparison of these findings with previously published stability studies(23).

## Methods

### Ethical approval

Animal experiment approval was obtained by the Institutional Animal Care and Use Committee (IACUC) at Rocky Mountain Laboratories. All animal experiments were executed in an Association for Assessment and Accreditation of Laboratory Animal Care (AALAC)-approved facility, following the guidelines in NIH Guide for the Care and Use of Laboratory Animals, Animal Welfare Act, United States Department of Agriculture and United States Public Health Service Policy on Humane Care and Use of Laboratory Animals. The Institutional Biosafety Committee (IBC) approved work with MERS-CoV strains under BSL3 conditions. Sample inactivation was performed according to IBC-approved standard operating procedures.

### Viral stock propagation

MERS-CoV and SARS-CoV-2 strains were obtained from different collaborators and passaged once in VeroE6 cells in DMEM (Sigma Aldrich) supplemented with 2% fetal bovine serum (Fisher Scientific), 1 mM L-glutamine (Thermo Fischer), 50 U/ml penicillin (Thermo Fischer) and 50 μg/ml streptomycin (Thermo Fischer). SARS-CoV-2 strain nCoV-WA1-2020 (MN985325.1) was provided by CDC, Atlanta, USA. Virus propagation was performed in VeroE6 cells in DMEM supplemented with 2% fetal bovine serum, 1 mM L-glutamine, 50 U/mL penicillin and 50 μg/mL streptomycin. VeroE6 cells were maintained in DMEM supplemented with 10% fetal bovine serum, 1 mM L-glutamine, 50 U/mL penicillin and 50 μg/mL streptomycin. Virus stocks were clarified by centrifugation and frozen at −80°C. Virus titrations were performed by end-point titration in VeroE6 cells inoculated with tenfold serial dilutions of virus. Cytopathic effect was scored at D5 (MERS-CoV) or D6 (SARS-CoV-2) and TCID_50_ was calculated from four replicates by the Spearman-Karber method (24).

### Sequencing stocks

All experiments, through second-strand cDNA synthesis, were performed in a BSLII cabinet for safety considerations. MERS-CoV samples were treated with RiboZero H/M/R rRNA (Illumina, San Diego, CA) depletion mix following the manufacturer’s instructions. After Ampure RNACleanXP (Beckman Coulter, Brea, CA) purification, the enriched RNA was eluted and assessed on a BioAnalyzer RNA Pico Chip (Agilent Technologies, Santa Clara, CA). These samples were then used to prepare second-strand cDNA, following the Truseq Stranded mRNA Library Preparation Guide, Revision E., (Illumina, San Diego, CA). To remove any remaining positive-strand RNA, samples were treated with RiboShredder RNase Blend. After AMpure XP purification (Beckman Coulter, Brea, CA), samples were analyzed on a RNA Pico chip to confirm RNA removal and the ends were adenylated following manufacturer’s recommendations. Final libraries were visualized on a BioAnalyzer DNA1000 chip (Agilent Technologies, Santa Clara, CA) and quantified using KAPA Library Quant Kit (Illumina) Universal qPCR Mix (Kapa Biosystems, Wilmington, MA) on a CFX96 Real-Time System (BioRad, Hercules, CA). Libraries were pooled together in equimolar concentrations and sequenced on the MiSeq (Ilumina, Inc, San Diego, CA) using on-board cluster generation and 2 x 250 paired-end sequencing. The cluster density was at 454k/mm2 per lane resulting in 8.7 Million reads passing filter per run with an average 85% > Q30.

### Phylogenetics

All available MERS-CoV genome sequences were downloaded from GenBank and curated to remove sequences that were not independently sampled. The GenBank MERS-CoV sequences were aligned with the consensus sequences for MERS-CoV isolates used in this study using the MAFFT v. 7.388 plugin (25) in Geneious Prime. The phylogenetic tree was inferred using the maximum likelihood method under the GTR + gamma model of nucleotide substitution with 1000 bootstrap replicates implemented with PhyML version 3.3.20190321.

### Stability of MERS-CoV on surface and in aerosols

Four different surfaces were evaluated: polypropylene (ePlastics), AISI 304 alloy stainless steel (Metal Remnants), copper (99.9%) (Metal Remnants) and silver (99.9%) (Sigma-Aldrich). Discs with a radius of 15 mm were cut out, sterilized and placed in 24-well plates. Each disc received 50 µl of MERS-CoV at a titer of 10^5^ TCID_50_/mL. At appropriate times, 1 mL of DMEM was added to the well, aliquoted and stored at -80°C. All samples were titrated on VeroE6 cells.

Virus stability in aerosols was determined as described previously (26). Briefly, the collison nebulizer used to produce aerosols was loaded with 10^6.5^ TCID_50_/ml of MERS-CoV in DMEM containing 2% FBS. Aerosols were maintained in the Goldberg drum and samples were collected at 0-, 30-, 60-, 120- and 180-minutes post aerosolization by passing air at 6L/min for 30 seconds from the drum through a 47mm gelatin filter (Sartorius). Filters were dissolved in 10 mL of DMEM containing 10% FBS and stored at -80°C. All samples were titrated on VeroE6 cells.

### Replication of MERS-CoV strains in vitro

VeroE6 cells were plated in 6-well plates and inoculated with an MOI of 0.01. Supernatant samples were obtained at 8, 24, 48 and 72 hours post infection (h.p.i.). Human airway epithelium inserts (HAE, Epithelix) were maintained as specified by manufacturer. HAEs were washed with 200 µl of phosphate-buffered saline for 30 minutes, followed by inoculation with MERS-CoV at an MOI of 0.1. Samples were obtained at 8, 24, 48, 72, and 96 h.p.i.

### Animal experiments

Transgenic *balb/c* mice expressing human DPP4 were inoculated intranasally (I.N.) with 10^3^ TCID_50_ MERS-CoV. Mice were weighed and oropharyngeal swabs were taken daily. At D3, four mice were euthanized, and lung tissue was harvested. The remaining six mice were monitored for survival. Mice were euthanized upon presence of severe disease signs (e.g. hunched posture, lack of movement) or >20% of weight loss.

### RNA extraction and quantitative reverse-transcription polymerase chain reaction

Tissues were homogenized and RNA was extracted using the RNeasy method (Qiagen) according to the manufacturer’s instructions. Swabs were added to 1 mL of DMEM, vortexed, and 140 µl was utilized for RNA extraction using the QiaAmp Viral RNA kit on the QIAxtractor. MERS-CoV viral RNA was detected via the UpE MERS-CoV assay (27) using the Rotor-GeneTM probe kit (Qiagen). MERS-CoV dilutions with known genome copies were run in parallel to allow calculation of genome copies in samples.

### Histology and immunohistochemistry

Necropsies and tissue sampling were performed according to IBC-approved protocols. Lungs were perfused with 10% formalin and processed for histologic review. Harvested tissues were fixed for a minimum of seven days in 10% neutral-buffered formalin and then embedded in paraffin. Tissues were processed using a VIP-6 Tissue Tek, (Sakura Finetek, USA) tissue processor and embedded in Ultraffin paraffin polymer (Cancer Diagnostics, Durham, NC). Samples were sectioned at 5 µm, and resulting slides were stained with hematoxylin and eosin. Specific anti-CoV immunoreactivity was detected using MERS-CoV nucleocapsid protein rabbit antibody (Sino Biological Inc.) at a 1:4000. The tissues were processed for immunohistochemistry using the Discovery ULTRA automated IHC/ISH staining instrument (Ventana Medical Systems) with a Discovery ChromoMap DAB (Ventana Medical Systems) kit, scanned with the Aperio ScanScope AT2 (Aperio Technologies, Inc.) and the entire section analyzed with the ImageScope Positive Pixel Count algorithm (version 9.1). All tissue slides were evaluated by a board-certified veterinary anatomic pathologist.

### Statistical analyses

All analyses were done using GraphPad Prism version 7.05 for Windows. All strains were compared to EMC/12. Linear regression was determined for the mean value of three runs per virus. Statistical significance was determined using ordinary one-way ANOVA followed by Bonferonni’s multiple comparisons test or a two-way unpaired student’s t-test.

## Results

### Stability of different MERS-CoV strains in aerosols or as fomites comparted to SARS-CoV-2

We utilized eight different MERS-CoV strains and one SARS-CoV-2 strain (SARS-CoV-2 strain nCoV-WA1-2020 (MN985325.1)) in the current study. Five MERS-CoV strains were isolated from human cases and three strains were isolated from dromedary camels. Strains were isolated between 2012 and 2018, and originated from the Middle East (5), Africa (2) or South Korea (1) (*Table 1*). All originally obtained viruses were passaged once in VeroE6 cells. Virus stocks were sequenced on the MiSeq. Mutations compared to the published sequence are detailed in Table 1.

**Table 1.** Details of MERS-CoV strains used in current study.

| Name | Host | Year | Location | Full name | Access number | SNPs |
| --- | --- | --- | --- | --- | --- | --- |
| **EMC/12** | Human | 2012 | KSA | HCoV-EMC/2012 | JX869059 | C6172T; C24059T; C24499A; G27162A |
| **U/14** | Human | 2014 | USA | Hu/Florida/USA-2/Saudi Arabia/2014 | KP223131 | None |
| **KSA/15** | Human | 2015 | KSA | Hu/Hofuf/KSA-11002/2015 | KY688120 | None |
| **SK/15** | Human | 2015 | South Korea | Hu/Korea/Seoul/177-3/2015 | KX034100 | C2149A; A6884G; T9566C; G10155T; A11376T; C14162T; C23041T; C26189T |
| **KSA/18** | Human | 2018 | KSA | Hu/Saudi Arabia/3015600912/2018 | MN723544 | C21149A; G22366A; C25009T |
| **C/KSA/13** | Camel | 2013 | KSA | Camel/Saudi Arabia/KFU-HKU1/2013 | KJ650297 | C25207T; C27875T |
| **C/E/13** | Camel | 2013 | Egypt | Camel/Egypt/NRCE/HKU270/2013 | KJ477103 | T16306C; C24100T; 26880T |
| **C/BF/15** | Camel | 2015 | Burkina Faso | Camel/Burkina Faso/CIRAD-HKU785/2015 | MG923471 | M29358C |

Available full-length MERS-CoV sequences were downloaded from Genbank. A phylogenetic maximum likelihood tree was constructed of the GenBank MERS-CoV sequences and the consensus sequences for the MERS-CoV isolates. The investigated MERS-CoV strains were distributed throughout the phylogenetic tree (*Figure 1*) and thus represent a broad sample of known genetic variation within currently circulating MERS-CoV strains.

**Figure 1.**
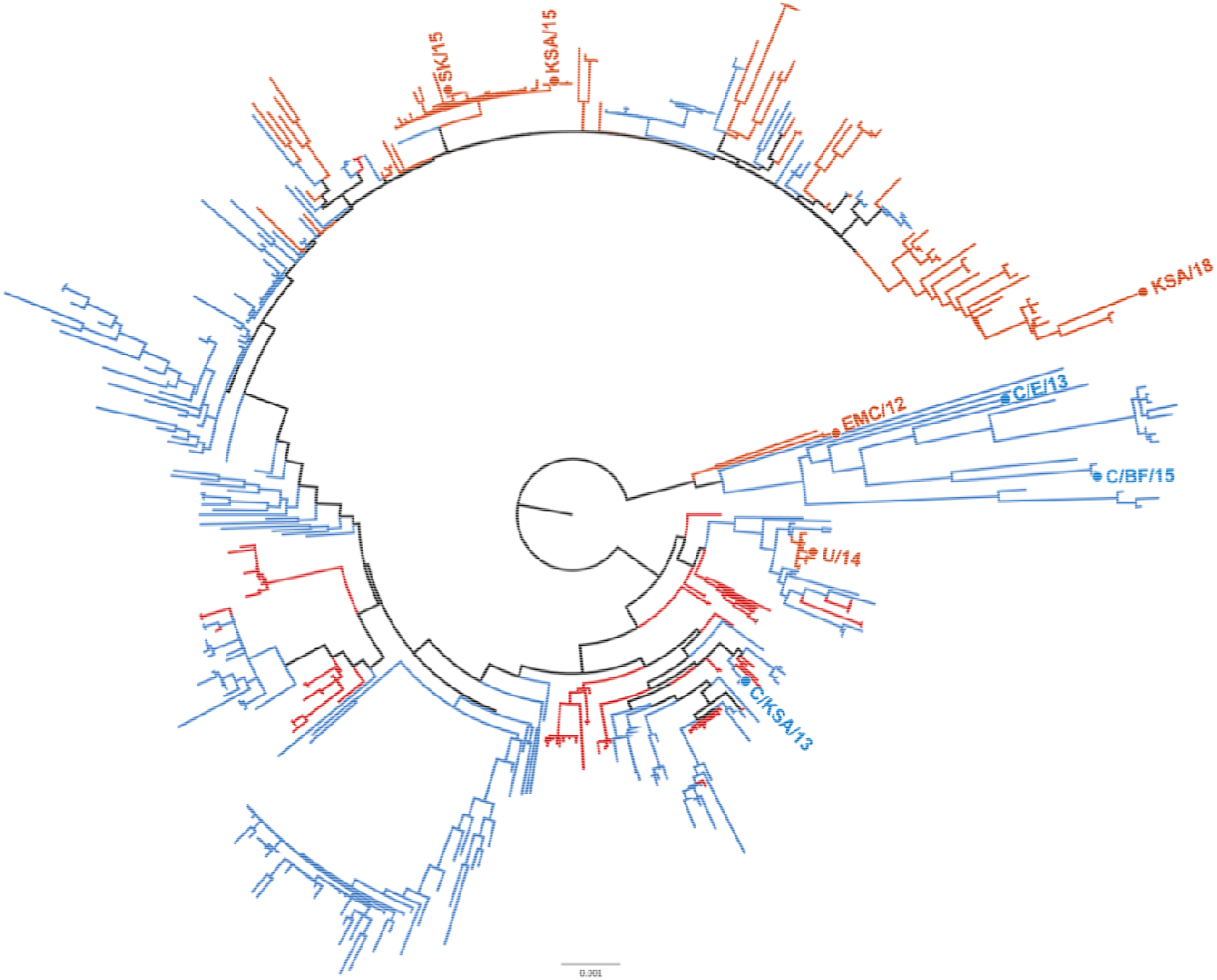
Phylogenetic tree of MERS-CoV strains. Maximum likelihood tree of 446 full MERS CoV genomes showing distribution of isolates used in this study. Human-derived MERS-CoV isolates used in this study are highlighted in red, camel-derived MERS-CoV isolates are highlighted in blue. Phylogenetic tree reconstructed with PhyML and rooted at the midpoint.

Stability of MERS-CoV strains was determined in aerosols as well as in fomites and compared to SARS-CoV-2. We investigated the stability of MERS-CoV as fomites on four different surfaces: polypropylene, stainless-steel, copper and silver. These surfaces were chosen as they represent commonly encountered surfaces in hospital environments or could function as a virocidal.

Back-titrations of all virus strains showed comparable starting virus titers. Stability of MERS-CoV on polypropylene and stainless-steel surfaces was similar to results previously reported for MERS-CoV and SARS-CoV-2 stability on surfaces (23, 28), except for strain C/KSA/13. Infectious virus titers of C/KSA/13 were significantly lower compared to EMC/12 on polypropylene (0, 1, and 24hrs) and stainless-steel surfaces (0, 24, and 48hrs). In contrast, infectious virus titers were low for all strains on copper and silver surfaces at 24hrs. Linear regression was calculated for the first 24hrs for each surface, and loss of infectious virus was significantly higher on copper and silver surfaces than on polypropylene and stainless-steel surfaces (−0.11576, −0.08744, −0.0529, and −0.0469 respectively) (*Figure 2A*).

**Figure 2.**
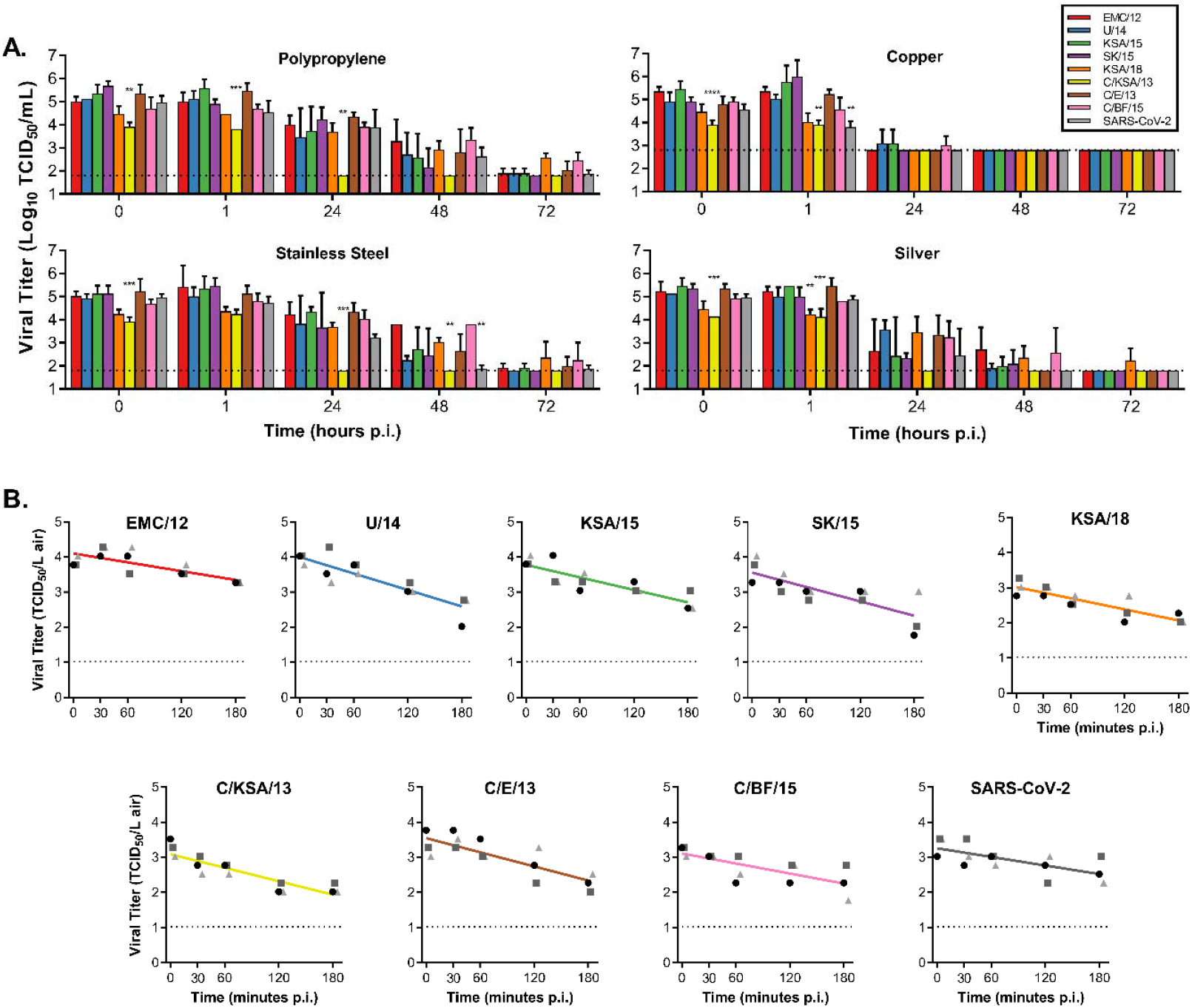
Stability of MERS-CoV strains on surfaces and in aerosols compared to SARS-CoV-2. A.) 50 µl of MERS-CoV or SARS-CoV-2 was spread on surface, either polypropylene, stainless steel, copper or silver. 1 mL of DMEM was added at T=0, 1, 24, 48 or 72 hours and titrated. B.) MERS-CoV or SARS-CoV-2 containing aerosols were sprayed into the Goldberg drum, and samples were taken at T=0, 30, 60, 120 and 180 minutes and titrated. Linear regression was calculated per virus and displayed in the graph as a line. A-B.) Statistically significant differences between EMC/12 and other strains were calculated using an unpaired Student’s two-tailed t-test corrected for multiple comparisons via Bonferroni. Dotted line = limit of detection; p-values = *<0.05; **<0.01, ***<0.001

All MERS-CoV strains were aerosolized in a Goldberg drum, samples were taken at 0, 30, 60, 120 and 180 min post aerosolization and titrated and compared with SARS-CoV-2. No significant differences in linear regression of loss of infectious virus in aerosols was detected between strains (EMC/12 = −0.00419; U/14 = −0.00781; KSA/15 = −0.00594; SK/15 = −0.00682; KSA/18 = −0.00527; C/KSA/13 = −0.00639; C/E/13 = - 0.00671; C/BF/15 = −0.00477). For all MERS-CoV strains, infectious virus could still be detected at 180 minutes post aerosolization (*Figure 2B*).

### In vitro replication of different MERS-CoV strains

Growth of all strains was then compared in two different *in vitro* cell systems: VeroE6 cells and HAE cultures. All strains were compared to the reference strain EMC/12 using a two-tailed unpaired Student’s t-test. At 48 hpi, C/KSA/13 and KSA/15 grew to significantly higher titers than EMC/12 in VeroE6 cells. At 72 hpi, C/KSA/13 and C/BF/15 grew to significantly lower titers than EMC/12 in HAE cultures. No other significant differences were observed in either VeroE6 cells or HAE cultures. While not always statistically significant, all camel-derived viruses had reduced replication kinetics as compared to EMC/12 in HAE cells at 24-72 hpi (*Figure 3*).

**Figure 3.**
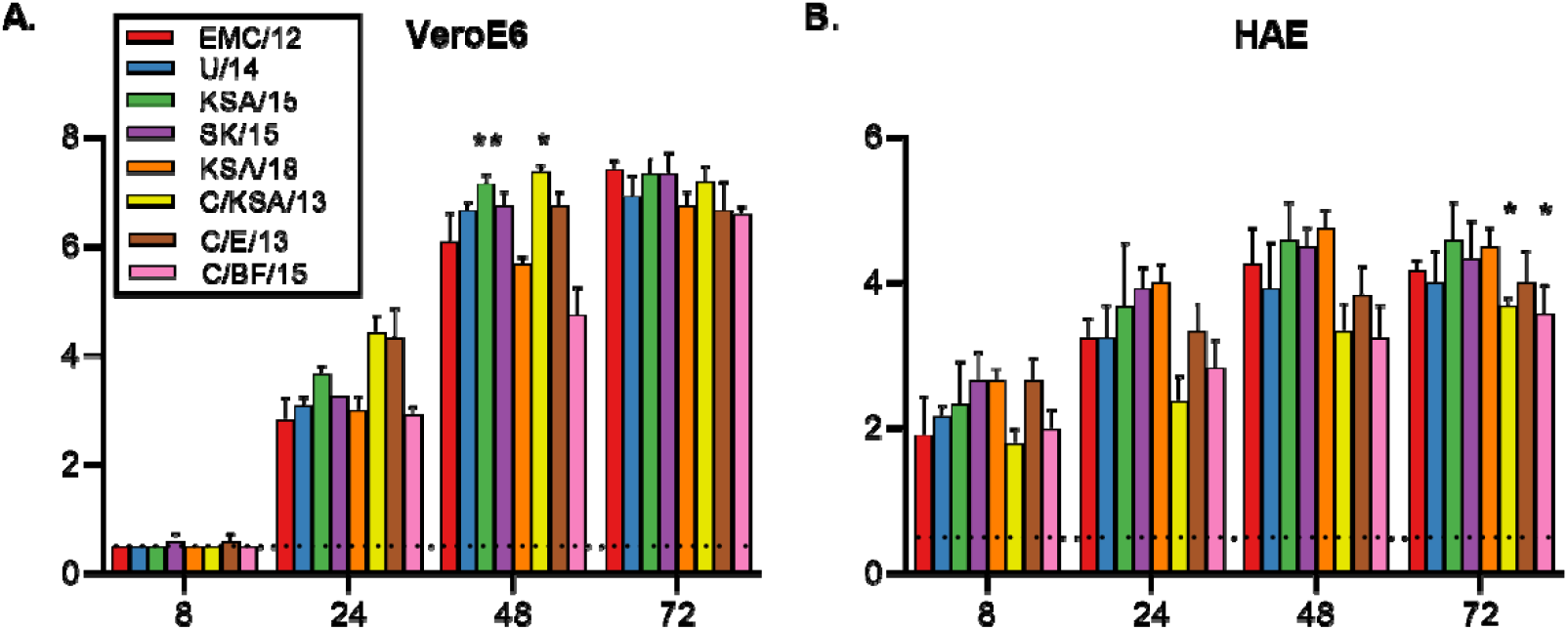
Virus replication in VeroE6 and human airway epithelium. Vero E6 cells (A) or HAE cultures (B) were infected with an MOI of 0.01 or 0.1 respectively, and samples of supernatant were obtained at 8, 24, 48 and 72 h.p.i. and titrated. Statistically significant differences as compared to the prototypical strain, EMC12, were calculated using an unpaired two-tailed Student’s t-test. Dotted line = limit of detection; p-values = *<0.05; **<0.01, ***<0.001

### Disease progression of different MERS-CoV strains in hDPP4 transgenic mice

MERS-CoV enters cells expressing the receptor human dipeptidyl peptidase IV (hDPP4). Our lab previously developed transgenic mice expressing hDPP4 to test MERS-CoV vaccine efficacy (22). Ten mice per group were inoculated I.N. with 10^3^TCID_50_ MERS-CoV per mouse. Mice started to lose weight on D2 to D5. Body weight kept decreasing for all groups, except for the mice inoculated with C/BF/15: only one mouse continued to lose weight (*Figure 4A*). This was accompanied by similar signs of disease across all groups; ruffled coat, increased breathing rate, reluctance to move, and hunched posture. No such signs were observed for mice inoculated with C/BF/15 that did not lose weight. Survivors were only found in the group inoculated with SK/15 (1 out of 6) and the group inoculated with C/BF/15 (5 out of 6). Average time to death was similar for all groups, excluding C/BF/15 (EMC/12 = 7.33; U/14 = 6.5; KSA/15 = 7; SK/15 = 7.6; KSA/18 = 7.67; C/KSA/13 = 7.5; C/E/13 = 8) (*Figure 4B*).

**Figure 4.**
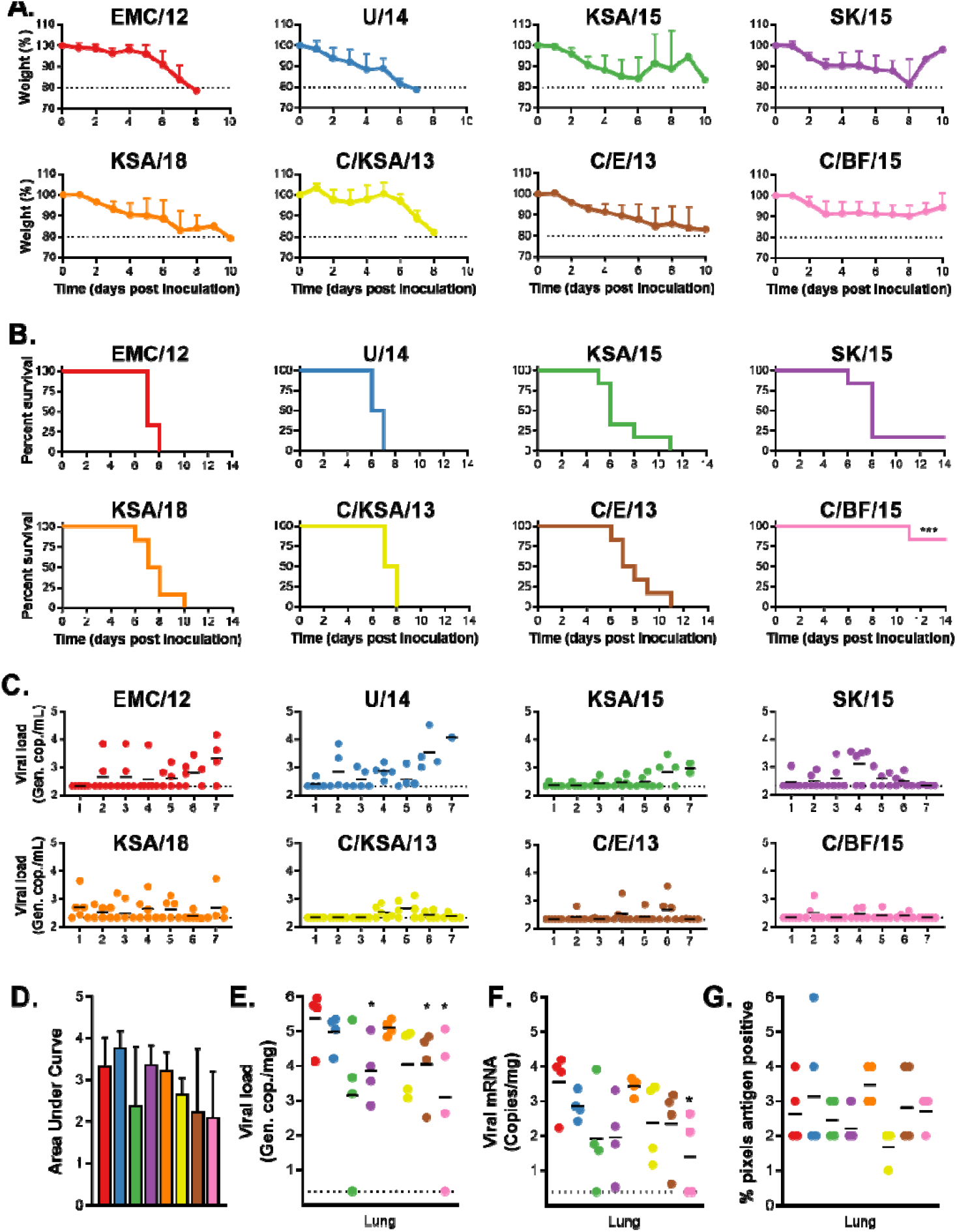
In vivo replication of different MERS-CoV strains. hDPP4 mice were inoculated I.N. with 10^3^ TCID_50_ MERS-CoV. Four mice were euthanized on D3, and the remaining 6 mice were monitored for survival. A.) Relative weight loss of hDPP4 mice. B.) Survival of hDPP4 mice. C.) Oropharyngeal shedding of MERS-CoV as measured via UpE qRT-PCR. D.) Area under the curve of oropharyngeal MERS-CoV shedding per virus strain. E.) Viral load in lung tissue obtained from mice euthanized at D3. F.) Viral mRNA load in lung tissue obtained from mice euthanized at D3. G.) Lung tissue were stained for MERS-CoV antigen and % of positive pixels was quantified. Statistical significance was compared using an unpaired two-tailed Student’s t-test. P-value = *<0.05.

Oral swabs were taken at D1 to D7 post inoculation and viral RNA was measured via qRT-PCR. The total amount of viral shedding was determined per group. No significant differences were found in the amount of shedding between different groups (*Figure 4C-D*). Viral gRNA and sgRNA was then measured via qRT-PCR in lung tissue obtained from four animals per group at D3. gRNA was significantly lower in lung tissue of mice inoculated with SK/15, C/E/15 and C/BF/15. sgRNA was only significantly lower in lung tissue of mice inoculated with C/BF/15 *(Figure 4E-F*).

Lung pathology was then examined by a board-certified veterinary pathologist blinded to study group allocation. No differences in pathology were observed. Animals rarely showed pulmonary pathology at D3, however animals that had lesions showed only a minimal and random lymphocytic infiltrate. Immunohistochemistry detecting MERS-CoV antigen was expressed rarely or randomly scattered in pulmonary tissue type I and II pneumocytes and not located in areas of inflammation. Morphometric analysis of pulmonary tissue with immunoreactivity revealed no significant difference between groups (*Figure 4G*).

## Discussion

The respiratory nature of MERS-CoV, in combination with its high mortality rate and frequent spillover from dromedary camels, position this virus as a potential threat to global health. The ongoing MERS-CoV endemic in the Middle East and subsequent discovery of the virus in camel herds across Africa has resulted in a wealth of publicly-available genetic data for various viral strains and isolates. Critically, a small number of studies have shown that genetic variation in MERS-CoV can drastically effect viral phenotypes including replication kinetics and pathogenicity (21, 29, 30). These findings highlight the need for MERS-CoV surveillance and, importantly, the assessment of new strains as they are isolated for mutations that increase spread, transmission and pathogenicity. In this study, we assessed several viral phenotypes as they relate to public health, in an attempt to better inform public health policy making with regards to MERS- and other respiratory-borne, human CoVs such as SARS-CoV-2. We assembled and tested a panel of diverse MERS-CoV viral isolates. Because MERS-CoV frequently spills over into the human population, we chose to include both human- and camel-derived strains (*Table 1, Figure 1*).

Nosocomial spread is at the center of MERS-CoV outbreaks. Therefore, we first assessed the stability of the virus on various surface material types commonly found in hospitals (polypropylene plastic and stainless steel) as well as materials with potential antiviral and known antimicrobial properties (silver and copper) (31, 32). Regardless of the surface material tested, C/KSA/13 was the least stable over time and fell below detectable levels by 24 hours (*Figure 2A*). As shown previously(23), all virus strains tested had notably reduced stability when left on copper. This finding was repeated on silver surfaces, with the copper surface proving most effective at reducing viral titers (*Figure 2A, right panels*). While copper and silver are generally appreciated for their antibacterial properties, copper has recently been shown to also have antiviral properties against influenza A H1N1 and SARS-CoV-2 (33-35). The exact mechanism of copper’s antiviral properties is still unclear, but may be related to the formation of hydroxyl radicals by copper ions when in aqueous solution (35). Silver-based nanoparticles have been shown to be antiviral for human immunodeficiency virus-1 (36), herpes simplex virus 2 (37), hepatitis B virus (38), respiratory syncytial virus (39), and monkey pox virus (40). Regardless of the mechanism, taking advantage of the antiviral properties of copper and silver could be a relatively straightforward method to decrease nosocomial transmission. Indeed, both silver and copper can be used for coating medical tools (41), and commonly touched items such as bed rails, door handles and intravenous poles (42). These findings appear to be more broadly applicable for other coronaviruses, as we observed similar results for SARS-CoV-2 (figure 2) (23). Further research should be invested in determining coronavirus susceptibility to copper-mediated inactivation.

MERS-CoV infects the lower respiratory tract in humans, and while the exact route of transmission has not been proven in a laboratory setting, it is likely to occur through aerosols and fomites (43). Studies have suggested that a hospital air-handling system may have contributed to nosocomial spread during the 2015 MERS-CoV outbreak in South Korea (16, 43) and our group has previously shown that the virus can remain viable suspended in air for up to 10 minutes (28). We tested aerosol stability over time for a diverse set of isolates with a Goldberg drum and observed that all viruses remained viable for a minimum of 180 minutes with approximately a log10 reduction in viral titer observed on average within the collected aerosols (*Figure 2B*). Even though we did not observe major differences in this study, strain stability is an important phenotype to continue monitoring, as mutations in viral capsid proteins have been shown to enhance environmental stability of bacteriophages, Dengue virus, and transmissible gastroenteritis virus (44-46). Because MERS-CoV isolates contain polymorphisms throughout the entire viral genome, including the structural proteins that form virions, it is still possible mutations may arise that influence overall virus particle stability. C/KSA/13, which showed reduced stability on surfaces in our experiments, contains polymorphisms in ORF1b, the spike glycoprotein, and the virion matrix protein in comparison to the other strains tested. Tracking and assessing the stability of coronavirus strains, even if to only rule out this possibility, will aid in our understanding of coronavirus variant spread during outbreaks, including the ongoing SARS-CoV-2 pandemic.

We tested viral replication kinetics of our virus panel in both VeroE6 cells as well as primary HAE cells (*Figure 3*). All virus isolates came up to similar titers on the VeroE6 cells by 72 hours, however KSA/15 and C/KSA/13 had a higher titer than EMC/12 by 48 hours post infection. Albeit not significant, C/BF/15 has a lower viral titer than EMC/12 at 48 and 72 hpi. These results are in good agreement with a previous study showing that C/BF/15 has impaired replication (20). In primary HAE cultures, all camel-derived viral isolates had reduced replication kinetics compared to EMC/12 (*Figure 3B*). More studies are needed with these camel-derived isolates to determine if their difference in replication kinetics results from a comparison with EMC12, which has well-described tissue culture adaptations, or to see if MERS-CoV may adapt in humans after transmission from camels.

Last, we tested our panel of viruses in a transgenic hDPP4 mouse model (22). We have previously shown that MERS-CoV replicates in type I and II pneumocytes within in the lower respiratory tract of this animal model (22). While MERS-CoV disease progression does not involve the central nervous system in humans, this small animal model is suitable for vaccine candidate testing, with animal survival or viral-induced death as a binary readout for vaccine efficacy. With the exception of C/BF/15 and SK/15, all strains tested were uniformly lethal in these animals, resulting in similar weight loss profiles and histopathology scores (*Figure 4*). MERS-CoV C/BF/15 contains a deletion in open reading frame 4b (ORF4b), which has been shown in a similar mouse model to result in impaired suppression of the host interferon response and increased type I and type III interferon signaling (20). Taken together, these results pave the way for testing MERS-CoV vaccine candidates for broadly neutralizing potential in this animal model (22, 47).

Our results with MERS-CoV C/KSA/13 suggest there may be a potential tradeoff between environmental surface stability and replication kinetics. Notably, this was observed for a camel-derived isolate, and we did not observe similar phenotypic relationships for the other strains tested (*Figure 2, 3*). Future research efforts with camel-derived viruses and more closely related human-derived viruses could reveal whether adaptations are likely to occur after zoonosis. Our previous viral stability results with SARS-CoV-2 and now these findings with MERS-CoV suggest copper should be incorporated more in hospital settings, particularly in areas of high contact between hospital workers and MERS patients, such as door handles, bed rails, and medical tools (23). Overall, we observed a range of stability, replication and pathogenesis phenotypes between different MERS-CoV isolates, underscoring the importance of continued surveillance of this virus and other coronaviruses such as SARS-CoV-2.

## Brief biography on the first authors

- 1. Dr. van Doremalen is a staff scientist in the Virus Ecology Section at the NIH’s Rocky Mountain Laboratories. Her research has helped develop animal models and test next generation vaccines for emerging infectious diseases.
- 2. Dr. Letko is an assistant professor at Washington State University. His laboratory studies how viral-host molecular interactions contribute to the potential for zoonotic transmission of novel, animal-derived viruses.

